# Mutational analysis reveals a novel role for hepatitis C virus NS5A domain I in cyclophilin-dependent genome replication

**DOI:** 10.1101/2023.07.18.549531

**Authors:** Shucheng Chen, Mark Harris

**Affiliations:** School of Molecular and Cellular Biology, Faculty of Biological Sciences and Astbury Centre for Structural Molecular Biology, University of Leeds, Leeds, LS2 9JT, United Kingdom

**Keywords:** hepatitis C virus, genotype, non-structural 5A protein (NS5A), genome replication, sub-genomic replicon, cyclophilin, cyclosporin

## Abstract

The hepatitis C virus (HCV) NS5A protein is comprised of three domains (D1-3). Previously, we observed that two alanine substitutions in D1 (V67A, P145A) abrogated replication of a genotype 2a (JFH-1 isolate) sub-genomic replicon (SGR) in Huh7 cells, but this phenotype was partially restored in Huh7.5 cells. To investigate the mechanism of this difference we extended this analysis to demonstrate that 5 additional residues, surface exposed and proximal to either V67 or P145, exhibited the same phenotype. In contrast, these mutants in a genotype 3a (DBN3a isolate) SGR retained their phenotype in each cell line.

The difference between Huh7 and Huh7.5 cells was reminiscent of the observation that cyclophilin (Cyp) inhibitors are more potent against HCV replication in the former and suggested a role for D1 in Cyp dependence. Consistent with this, all JFH-1 and DBN3a mutants exhibited increased sensitivity to cyclosporin A treatment compared to wildtype. Silencing of CypA in Huh7 cells inhibited replication of both JFH-1 and DBN3a. However, in Huh7.5 cells CypA silencing did not inhibit JFH-1 wildtype, but abrogated replication of all the JFH-1 mutants, and both DBN3a wildtype and all mutants. CypB silencing in Huh7 cells had no effect on DBN3a, but abrogated replication of JFH-1. CypB silencing in Huh7.5 cells had no effect on either SGR. These data demonstrate both a direct involvement of NS5A D1 in Cyp-dependent genome replication and functional differences between genotype 2 and 3 NS5A. Lastly, we confirmed that JFH-1 NS5A D1 interacted with CypA *in vitro*.

## Introduction

Hepatitis C virus (HCV) belongs to the *Hepacivirus* genus of the *Flaviviridae* family and is now formally named as the species *Hepacivirus hominis*. HCV infection is one of the leading causes of acute and chronic hepatitis, leading to chronic liver diseases such as liver cirrhosis and hepatocellular carcinoma (HCC). According to the WHO 58 million people are infected with HCV worldwide and nearly 300,000 deaths every year are attributed to this virus [1]. HCV is classified into 8 genotypes with an uneven geographical distribution. Notably genotype 3 is the second most prevalent worldwide and is associated with more severe pathology than other genotypes [2]. Genotype 3 infected patients are also most likely to exhibit resistance to direct acting antivirals (DAA), which are currently in widespread clinical use and are highly efficacious (over 95% sustained virological response (SVR)) [3].

HCV is enveloped with a positive-sense, single-stranded RNA of approximately 9.6 kb [4]. The genome contains a single large open reading frame encoding a 3000-residue polyprotein flanked by 5′ and 3′ untranslated regions. The polyprotein is cleaved by viral and host proteases into 3 structural (Core, E1 and E2), and 7 non-structural (p7, NS2, NS3, NS4A, NS4B, NS5A and NS5B) proteins. Of note, NS3 protease, NS5A and NS5B RNA-dependent RNA polymerase are the targets of DAAs.

NS5A is a phosphoprotein with multiple roles in the HCV life cycle. NS5A comprises an N-terminal amphipathic α-helix membrane anchor and three domains (D1, D2 and D3) which are connected by two solvent-exposed, low-complexity sequences (LCS-I and LCS-II) (Fig 1A). D1 is the only structured domain and is required for genome replication, in addition our recent data has shown that D1 is also involved in virus assembly [5], likely by blocking a novel function of PKR [6]. D2 is a disordered domain that has been reported to interact with cyclophilin A (CypA) [7, 8], a member of the Cyp family of peptidyl prolyl isomerases required for HCV genome replication [9]. D3 has also been reported to interact with CypA [10], and NS5B has been reported to be regulated by binding to the related CypB [11]. Cyps are one of the most abundant proteins in the cytoplasm (0.1–0.4% of total protein content) and are expressed in all tissues [12]. The mechanisms by which Cyps contribute to viral replication remain obscure, however they have been proposed to both induce conformational changes in target proteins (as a result of isomerase activity), and/or regulate protein complex formation [13]. An emerging theme is that Cyps protect either viral particles or replication complexes from cellular antiviral factors such as innate immune sensors [13]. Notably, both CypA and CypB are targets of the immunosuppressive drug cyclosporin A (CsA), and consequently CsA was demonstrated to inhibit HCV genome replication [14].

**Fig 1.**
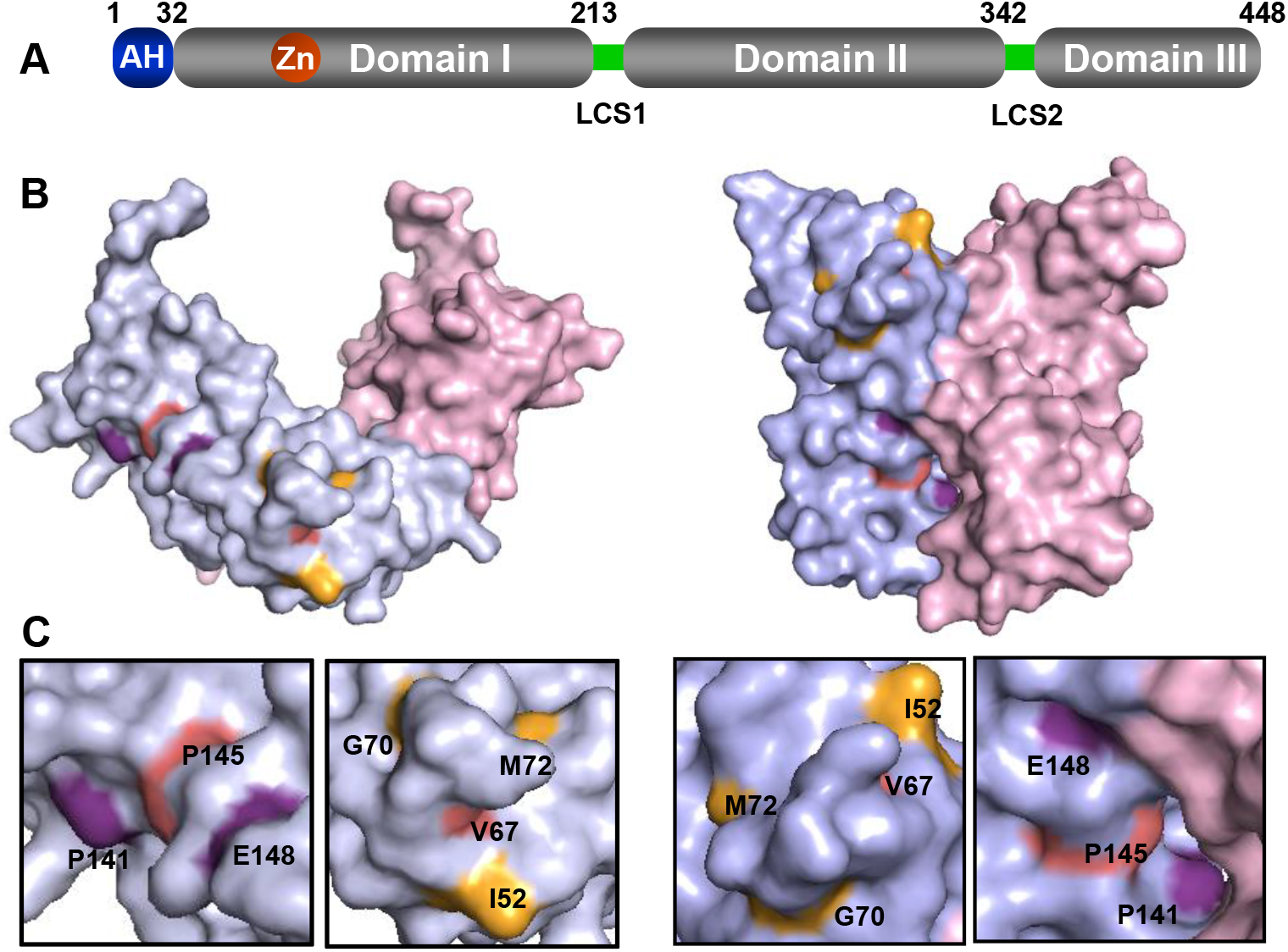
Structure of NS5A D1 and location of mutated residues. **(A)** Structure of NS5A illustrating the three domains. AH: amphipathic helix (purple), LCS: low complexity sequence (yellow). **(B, C)** Conserved and surface exposed residues proximal to V67 and P145 (red) are displayed in the two NS5A D1 (genotype 1b) structures 1ZH1 (**B)** and 3FQM **(C)**.

In this study, we use a combination of mutagenesis and silencing of Cyp expression to extend our understanding of the role of Cyps in HCV biology by showing that NS5A D1 also contributes to the dependence of HCV on Cyps for genome replication. We further show that two HCV genotypes (2a and 3a) exhibit differential requirements for CypA and CypB.

## Results

### Two clusters of surface exposed residues in NS5A D1 exhibit partial defects in replication

We previously showed that alanine substitution of residues V67 and P145 within NS5A D1 resulted in a partial defect in genome replication in Huh7 cells but exhibited no replication phenotype in Huh7.5 cells [5]. To further understand this phenotype we sought to identify additional residues which also exhibited the cell line specific replication requirements of V67 and P145. To achieve this goal, we interrogated the two genotype 1b monomeric structures of NS5A domain I (PDB 1ZH1 and 3FQM) and identified 15 surface exposed and conserved residues proximal to V67 and P145 [6]. These residues were targeted for alanine scanning mutagenesis in the context of an SGR containing a firefly luciferase reporter and unique restriction sites engineered either side of the NS5A coding sequence to facilitate sub-cloning (mSGR-luc-JFH-1) [15, 16]. Analysis of the replication of this panel of NS5A D1 mutant SGRs in both Huh7 and Huh7.5 cell lines revealed several distinct phenotypes ranging from no effect to complete abrogation of replication in both cell lines [6]. Here we focus our attention on five residues that exhibited the same genome replication phenotype as V67A and P145A, ie a partial and significant defect in Huh7 cells which was restored to wildtype levels in Huh7.5 cells (Fig 2). Three of these residues were proximal to V67 (I52, G70 and M72), and two were proximal to P145 (P141 and E148) (Figs 1B and 1C).

**Fig 2.**
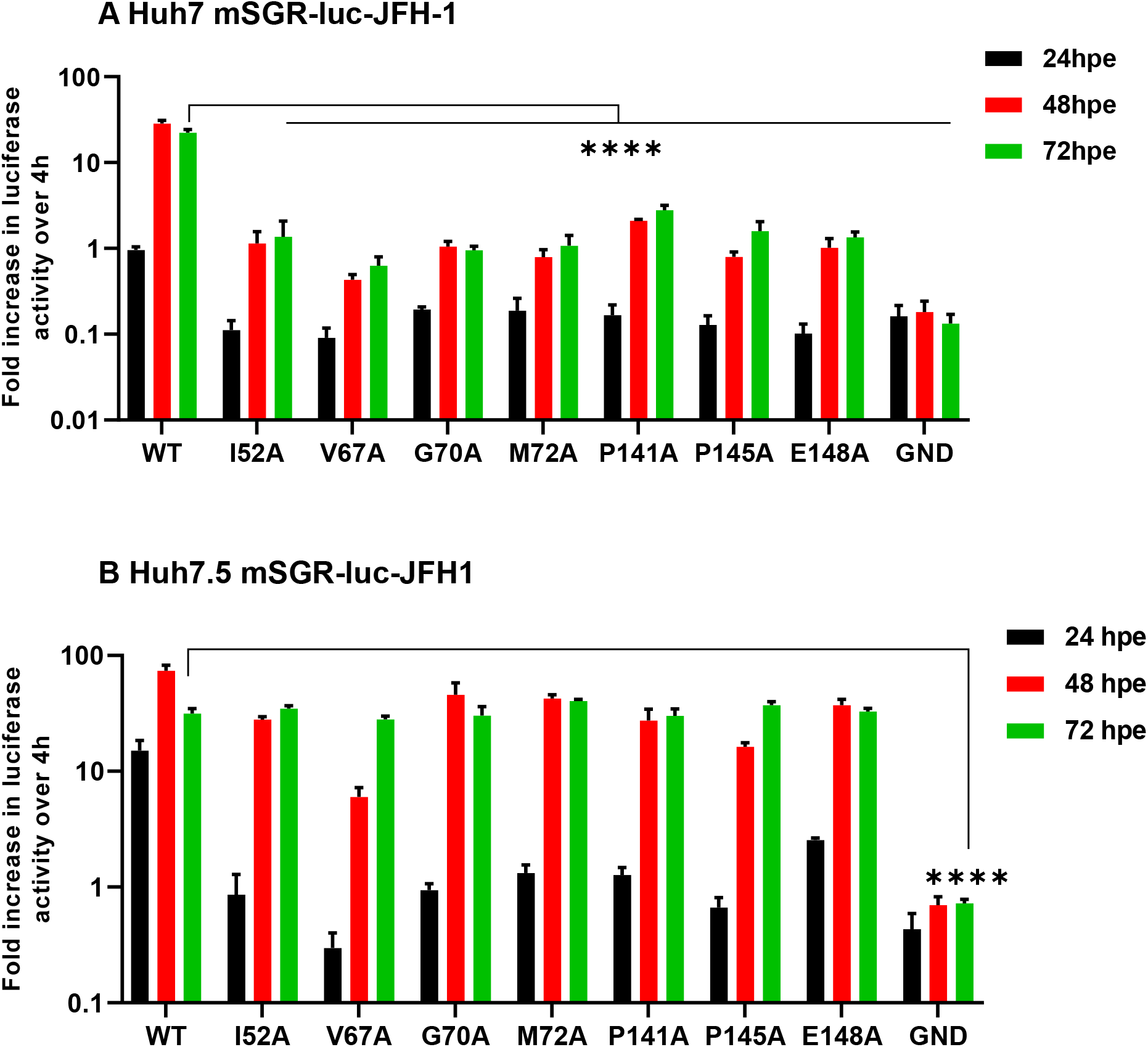
Genome replication phenotypes of JFH-1 mutants in Huh7 and Huh7.5 cells. *In vitro* transcribed mSGR-luc-JFH-1 RNAs containing the indicated mutations were electroporated into **(A)** Huh7 and **(B)** Huh7.5 cells. Firefly luciferase activity was measured at 4, 24, 48 and 72 hpe and the data were normalized with respect to 4 hpe. N=3, significant differences from WT denoted by **** (P<0.0001).

### D1 mutant replication phenotypes are not conserved in the context of a genotype 3a SGR

The majority of molecular studies of HCV have been performed using the genotype 2a JFH-1 isolate, however recently infectious clones have been developed for other genotypes. Of particular interest is genotype 3 as this is associated with more rapid fibrosis progression, a higher incidence of steatosis and hepatocellular carcinoma, and lower response rates to DAA treatment. As part of a broader programme of work to compare the functions of NS5A between different genotypes, we extended this current analysis to the genotype 3a isolate, DBN3a [17]. We recently developed an SGR derived from the DBN3a infectious clone [18] termed SGR-luc-DBN3a, and we generated the same 7 alanine substitutions as described in Fig 2 for JFH-1. Note that all of these residues are conserved apart from I52 which is V52 in DBN3a (Supp. Fig S1).

Analysis of the replication of the SGR-luc-DBN3a mutant panel as shown in Fig 3 revealed significant differences from JFH-1. With the exception of P141A which was indistinguishable from wildtype in both cell lines, the other mutants all showed replication defects that were more pronounced in Huh7 than Huh7.5 cells. However, unlike JFH-1, only V52A out of these 6 mutants was restored to wildtype levels of replication in Huh7.5 cells (Fig 3B). In particular P145A exhibited the lowest levels of replication, and unlike JFH-1, this mutant in DBN3a was actually less active in Huh7.5 cells. These data point to fundamental differences in the functions of JFH-1 and DBN3a NS5A.

**Fig 3.**
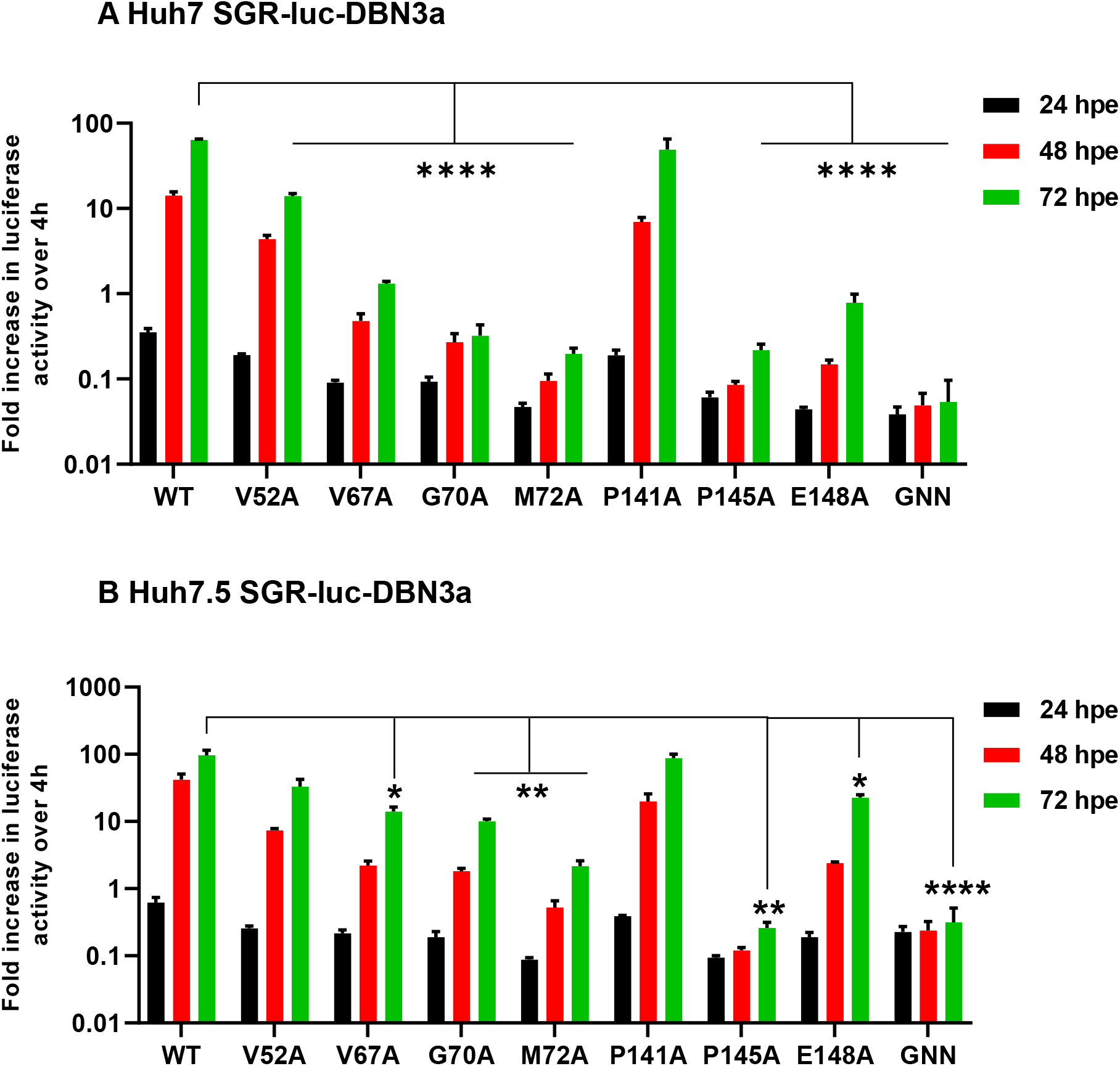
Genome replication phenotypes of DBN3a mutants in Huh7 and Huh7.5 cells. *In vitro* transcribed SGR-luc-DBN3a RNAs containing the indicated mutations were electroporated into **(A)** Huh7 and **(B)** Huh7.5 cells. Firefly luciferase activity was measured at 4, 24, 48 and 72 hpe and the data were normalized with respect to 4 hpe. N=3, significant differences from WT denoted by * (P<0.05), ** (P<0.01) and **** (P<0.0001).

### D1 mutants exhibit increased sensitivity to cyclosporin A (CsA) treatment

The differences that we observed between the two cell lines, in particular for JFH-1, were reminiscent of a study published recently [19] demonstrating that cyclophilin (Cyp) inhibitors were more potent against HCV replication in Huh7 compared to Huh7.5 cells. We reasoned that our data might point to a role for D1 in Cyp dependence. To address this we determined the CsA sensitivity of the JFH-1 mutant panel in Huh7.5 cells - we chose these cells as all the mutants were able to replicate efficiently in these cells, and in contrast the impaired replication in Huh7 cells might confound interpretation of the data. As a control we used the well characterized D2 mutant D316E which was previously shown to be highly resistant to CsA [8]. As expected, D316E showed a 7-fold increase in CsA EC_50_ over wildtype (Fig 4), in contrast all of the D1 mutants were more sensitive to CsA with significantly reduced EC_50_ values compared to wildtype. The same was true for DBN3a, with all mutants that could replicate in Huh7.5 cells showing a reduced CsA EC_50_ value compared to wildtype (Fig 5). To confirm that the increased sensitivity to CsA was not due to reduced replicative capacity we plotted the EC_50_ value against the replication level (as measured by the ratio of firefly luciferase between 48/4 h). Although there was a trend towards increased CsA sensitivity with lower replication, there were significant outliers for both JFH-1 and DBN3a. We conclude that for both JFH-1 and DBN3a, D1 contributes to CsA sensitivity implying that D1 may be involved in interactions with Cyps that contribute to genome replication.

**Fig 4.**
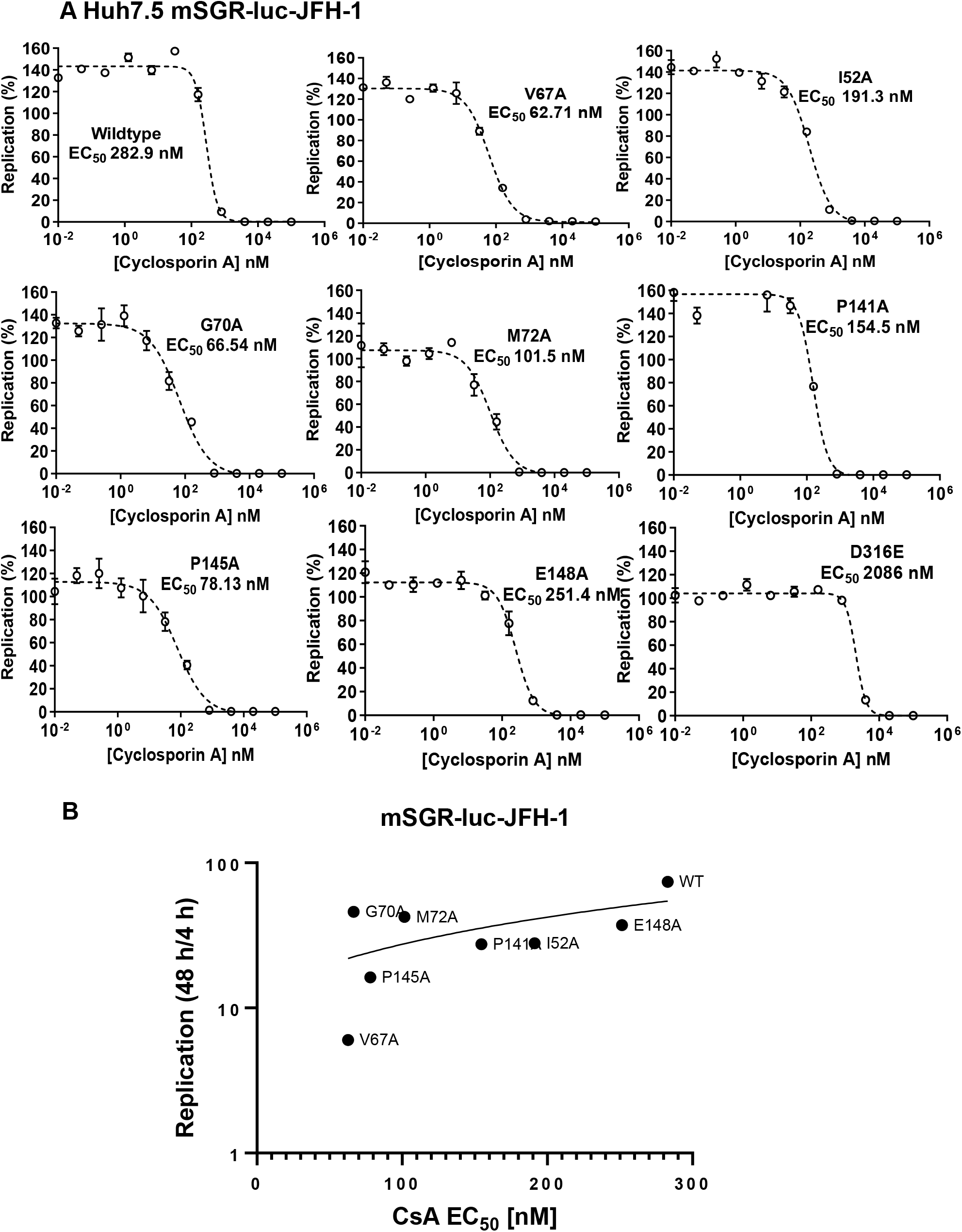
CsA sensitivity analysis of JFH-1 NS5A D1 mutants. **(A)** Huh7.5 cells were electroporated with mSGR-luc-JFH-1 RNAs containing the indicated mutations, seeded into 96-well plates and treated with 5-fold serial dilutions of CsA. Cells were harvested at 48 hpe and firefly luciferase activity measured. Data were normalized to the DMSO control and processed using the EC_50_ model of GraphPad 9.30. **(B)** CsA EC_50_ values were plotted against replicative capacity of each mutant (as determined by the ratio of firefly luciferase at 48 hpe to 4 hpe).

**Fig 5.**
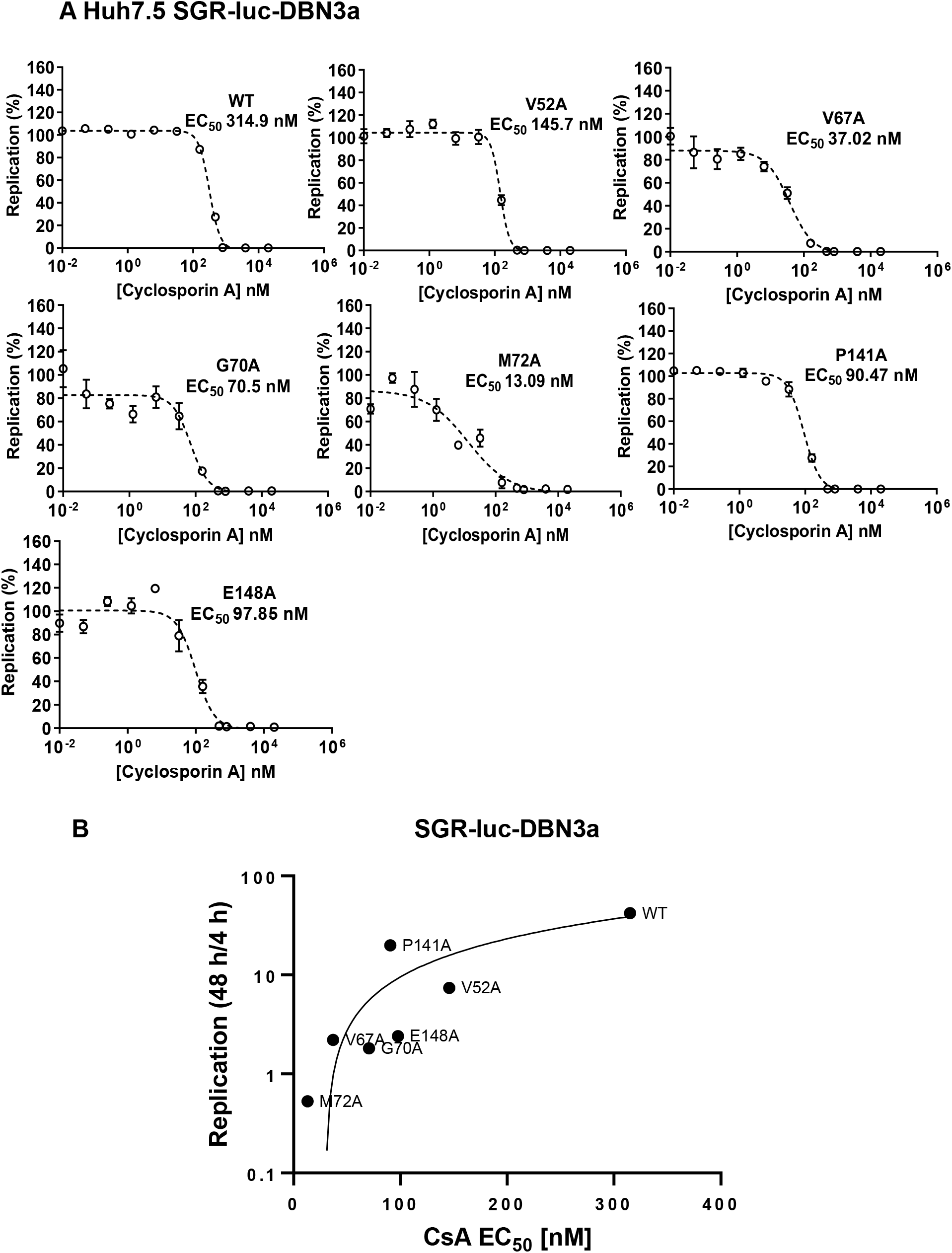
CsA sensitivity analysis of DBN3a NS5A D1 mutants. **(A)** Huh7.5 cells were electroporated with SGR-luc-DBN3a RNAs containing the indicated mutations, seeded into 96-well plates and treated with 5-fold serial dilutions of CsA. Cells were harvested at 48 hpe and firefly luciferase activity measured. Data were normalized to the DMSO control and processed using the EC_50_ model of GraphPad 9.30. **(B)** CsA EC_50_ values were plotted against replicative capacity of each mutant (as determined by the ratio of firefly luciferase at 48 hpe to 4 hpe).

### D1 mutants reveal differential cyclophilin dependence of JFH-1 and DBN3a genome replication

CsA binds to and inhibits Cyps, although 18 of these enzymes have been identified in humans only two (CypA and CypB) have been shown to be required for HCV genome replication [9, 11]. Given the changes in CsA sensitivity exhibited by the D1 mutants we therefore hypothesised that either CypA and/or CypB might regulate D1 function in genome replication. To test this we generated CypA and CypB silenced Huh7 and Huh7.5 cell lines using lentivirus delivery of shRNA, with a non-specific shRNA as control. Silencing was verified by western blotting (Figs 6A and B) and these cell lines were then electroporated with RNAs from the two panels of NS5A D1 mutant SGRs (JFH-1 and DBN3a).

**Fig 6.**
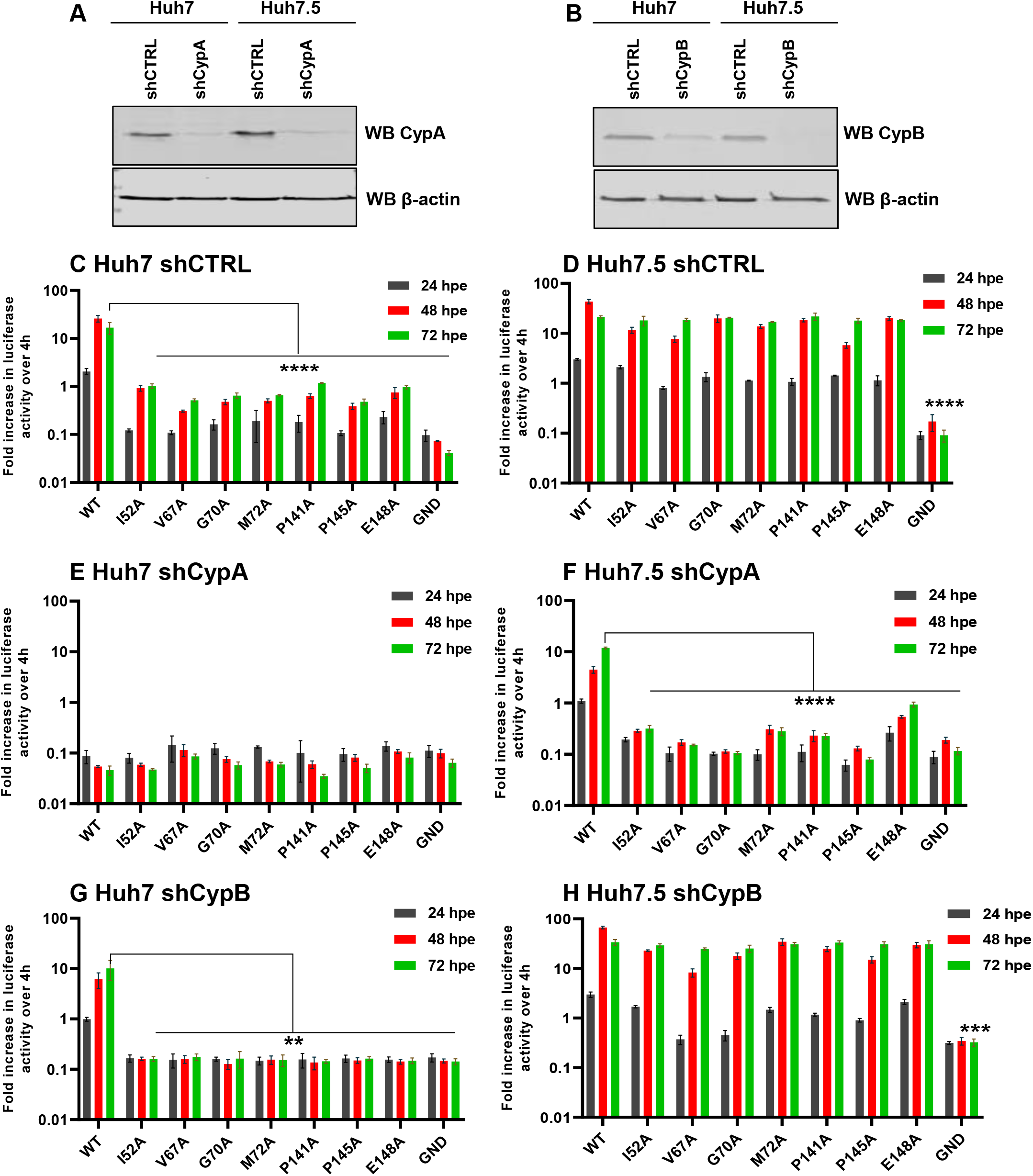
Genome replication phenotypes of JFH-1 mutants in Cyp silenced cells. **(A, B)** CypA **(A)** and Cyp B **(B)** expression was detected in silenced Huh7 and Huh7.5 cells by western blot. **(C-H)** *In vitro* transcribed mSGR-luc-JFH-1 RNAs containing the indicated mutations were electroporated into the indicated cell lines. Luciferase activity was measured at 4, 24, 48 and 72 hpe and the data were normalized with respect to 4 hpe. N=3, significant differences from WT denoted by ** (P<0.01), *** (P<0.001) and **** (P<0.0001).

Reassuringly, in the shCTRL cell lines the profile of replication for all SGRs mirrored that seen in parental Huh7 and Huh7.5 (compare Figs 2A with 6C/D, and 3A with 7A/B), confirming that lentivirus transduction and/or puromycin selection did not affect HCV replication. For both JFH-1 and DBN3a replication of both wildtype and the mutants was completely abolished by silencing of CypA in Huh7 cells (Figs 6E and 7C). In Huh7.5 CypA silenced cells JFH-1 wildtype retained the ability to replicate, albeit reduced compared to parental and shCTRL cells (Fig 6F). This was consistent with the previous observation that CypA is critical for HCV replication in Huh7 cells, but not in Huh7.5 cells [19]. However, none of the JFH-1 mutants were able to replicate in CypA-silenced Huh7.5 cells, with the exception of E148A which exhibited a very low level of activity (Fig 6F). In contrast, for DBN3a no replication was observed in CypA-silenced Huh7.5 cells for either wildtype or the mutants (Fig 7D). We conclude that DBN3a genome replication is absolutely dependent on CypA in comparison to JFH-1.

**Fig 7.**
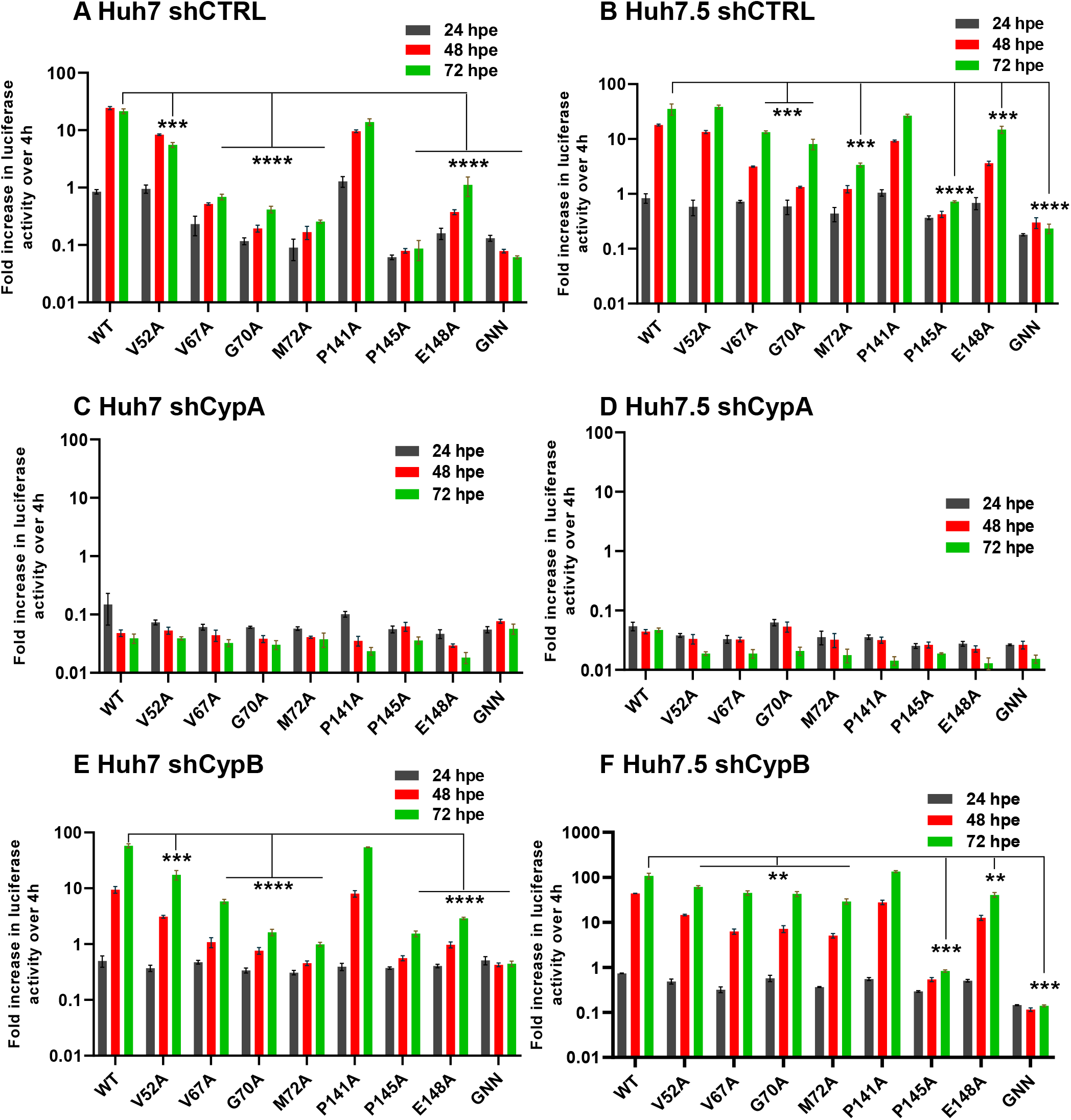
Genome replication phenotypes of DBN3a mutants in Cyp silenced cells. **(A-F)** *In vitro* transcribed SGR-luc-DBN3a RNAs containing the indicated mutations were electroporated into the indicated cell lines. Luciferase activity was measured at 4, 24, 48 and 72 hpe and the data were normalized with respect to 4 hpe. N=3, significant differences from WT denoted by ** (P<0.01), *** (P<0.001) and **** (P<0.0001).

The situation in CypB silenced cells was very different. For JFH-1 only the wildtype could replicate in CypB silenced Huh7 cells (Fig 6G), whereas in Huh7.5 cells CypB silencing had no effect, being comparable to parental or shCTRL cells (Fig 6H). In contrast, CypB silencing had no effect at all on DBN3a replication in either cell line (Fig 7E/F). Thus DBN3a exhibits no dependence on CypB compared to JFH-1.

### NS5A D1 interacts with CypA

Previously it has been demonstrated that CypA binds to a proline-rich region within NS5A D2, with prolines in D2 also acting as substrates for the peptidyl-prolyl isomerase activity of CypA [20]. As we observed that the panel of D1 mutants resulted in increased sensitivity of genome replication to CsA, and abrogated the ability of the wildtype JFH-1 SGR to replicate in CypA-silenced Huh7.5 cells, we proposed that CypA might also interact with D1. To test this, we performed an *in vitro* GST-pulldown assay using GST-CypA (wildtype or H126Q catalytically inactive mutant) as baits to precipitate purified His-tagged NS5A D1 (residues 35-215). As shown in Fig 8A, wildtype CypA did indeed precipitate NS5A D1, whereas the CypA-H126Q mutant was unable to do so suggesting that the interaction of D1 with CypA is dependent on the active isomerase site. To confirm this hypothesis we demonstrated that the CypA-NS5A D1 interaction was inhibited by the presence of CsA (Fig 8B). Lastly we expressed the JFH-1 D1 mutant I52A as an exemplar, and investigated the effect of this mutation on the interaction with CypA. As shown in Fig 8C, I52A was not precipitated by GST-CypA (wildtype or H126Q). We conclude from these data that NS5A D1 is able to physically interact with CypA and this interaction was dependent on the peptidyl-prolyl isomerase actvitiy of CypA. This interaction contributes to the CypA dependence of HCV genome replication.

**Fig 8.**
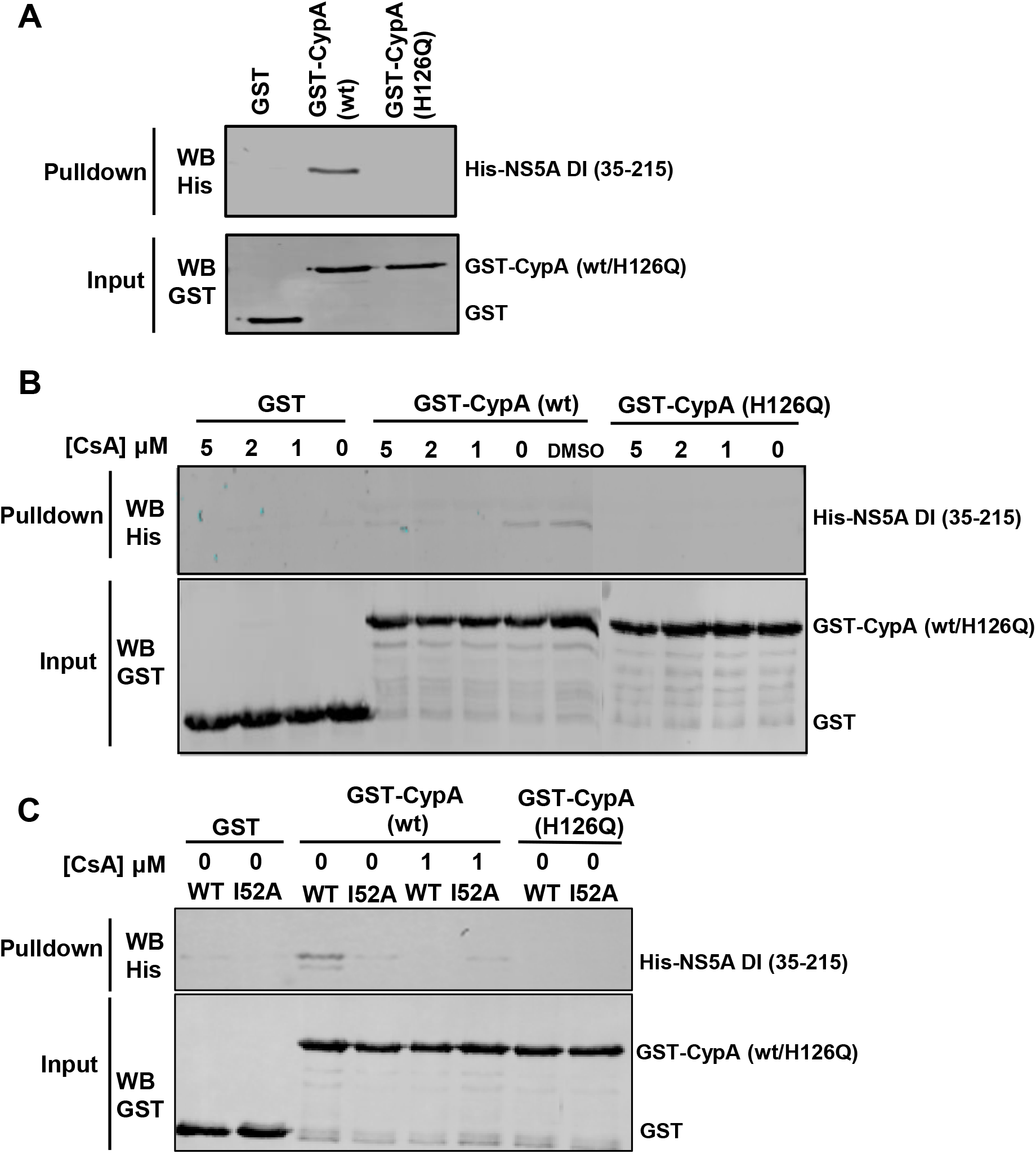
Interaction between JFH-1 NS5A D1 and CypA. **(A)** GST, GST-CypA (wildtype) and GST-CypA (H126Q) were purified and bound to glutathione-agarose as a bait to precipitate purified His-SUMO tagged JFH-1 NS5A D1 (amino acids 35-215). The precipitates were analysed by western blotting for the His-tag (top), and inputs verified by GST western blotting (bottom). **(B)** Indicated concentrations of CsA were added to GST or GST-CypA loaded glutathione-agarose for 10 min before the addition of His-SUMO tagged JFH-1 NS5A D1 and analysis as in (A). **(C)** His-SUMO tagged JFH-1 NS5A D1 (wildtype or I52A mutant) were added to GST or GST-CypA loaded glutathione-agarose and analysed as in (A) and (B). Where indicated, CsA was added prior to addition of the NS5A D1 prey.

## Discussion

The data presented in this paper attest to significant differences in the functions of NS5A D1 from JFH-1 and DBN3a. This is perhaps surprising given that the D1 amino acid sequence is 72% identical between the two isolates (Supp. Fig S1). However, it is well documented that genotype 3 HCV is associated with more severe disease progression and higher resistance to DAAs, particularly those targeting NS5A D1 [3]. Thus it is possible that these characteristics are driven by subtle differences in protein structure and/or interactions with cellular factors. In this regard, our data also show that JFH-1 and DBN3a exhibit a differential requirement for CypA and CypB. Specifically, DBN3a is more dependent on CypA than JFH-1, as wildtype DBN3a failed to replicate in either Huh7 or Huh7.5 in which CypA was silenced, whereas wildtype JFH-1 was able to replicate in CypA silenced Huh7.5 (Figs 5F and 6D). This suggests that JFH-1 NS5A D1 binds a cellular antiviral factor that in the absence of CypA blocks HCV genome replication, whereas DBN3a D1 is unable to target this factor, rendering it more sensitive to CypA silencing. It is tempting to speculate that this might in part explain the different kinetics of DBN3a and JFH-1 genome replication, with DBN3a exhibiting a significant lag phase before replication proceeds (compare Figs 2 and 3). This hypothesis is supported by the observation that JFH-1 wildtype also exhibits a lag in CypA silenced Huh7.5 cells (Fig 6F).

In contrast, DBN3a did not exhibit any requirement for CypB (Fig 7 E/F) compared to JFH-1 for which only wildtype was able to replicate in CypB silenced Huh7 cells. CypB was initially reported to interact with NS5B and stimulate RNA-dependent RNA polymerase activity [11], however CypB has also been reported to interact with NS5A D2 [20]. Our data are consistent with the suggestion that CypB also targets NS5A D1 of JFH-1 but not that of DBN3a. Clearly the differential requirement of the two isolates for the two Cyps demonstrates that these two cellular factors are not redundant with each other, and likely function via distinct mechanisms to facilitate viral genome replication.

All mutants in D1 tested in this study increased the sensitivity to CsA (Figs 4 and 5). Furthermore, in the case of JFH-1 they blocked replication in CypA silenced Huh7.5 cells and CypB silenced Huh7 cells suggesting that the residues mutated may be involved in direct interactions with the two Cyps, at least for JFH-1. The direct interaction of D1 with CypA *in vitro*, and the loss of the CypA interaction exhibited by I52A (Fig 8), provides some supporting evidence for this hypothesis. The disruption of this interaction by CsA or a mutation in the CypA active site suggests that CypA might function to isomerise D1, perhaps at one of the two prolines targeted in the mutagenesis. A key question that arises from this study is the molecular basis underpinning the differences in Huh7 and Huh7.5 cells? One possibility is that the antiviral effector PKR is proposed to be regulated by CypA [19] and was reported to be expressed at lower levels in Huh7.5. However, in our hands we see very little effect on genome replication when PKR is silenced in either Huh7 or Huh7.5 cells (Supp. Fig. S2), although in other work we have shown a novel role for D1 in blocking PKR-mediated inhibition of virus assembly [6]. It is also documented that Huh7.5 are heterozygous for a dominant negative mutant of RIG-I [21]. Again, we do not believe that RIG-I inactivation is the explanation as either RIG-I silencing in Huh7 cells, or exogenous RIG-I over-expression in Huh7.5 cells, had no effect on JFH-1 SGR replication (Supp. Fig. S3). Further studies in our laboratory are in progress to dissect out the molecular differences between JFH-1 and DBN3a NS5A, as well as the pathways associated with CypA and PKR that regulate the virus lifecycle.

## Acknowledgements

This work was supported by a Wellcome Investigator Award (grant number 096670), and an MRC project grant (MR/S001026/1) to MH. SC was supported by a University of Leeds/China Scholarship Council PhD studentship. We thank Greg Towers (UCL) for the Cyp silencing lentivirus vector constructs. The funders had no role in study design, data collection and analysis, decision to publish, or preparation of the manuscript.

## Materials and Methods

### Cell lines

Huh7 (human hepatocellular carcinoma) [22] and Huh7.5 cells (a derivative of Huh7 from which a stable subgenomic replicon was ‘cured’ by IFN treatment and which exhibit a defect in RIG-I) [21, 23] were used for electroporation. Cells were cultured in Dulbecco’s Modified Eagles Medium (DMEM; Sigma) supplemented with 10 % foetal bovine serum (FBS), 100 IU penicillin/ml, 100 µg/ml streptomycin and 1 % non-essential amino acids (Lonza) in a humidified incubator at 37°C, 5 % CO_2_.

### Plasmid and virus constructs

The JFH-1 derived SGR with a luciferase reporter (mSGR-luc-JFH1) was described previously [16], and contains BamHI/AfeI restriction sites flanking the NS5A coding sequence. The DBN3a SGR was generated from the DBN3acc genotype 3a infectious clone [17], and was described recently [18]. NS5A mutations were constructed using Q5 Site-Directed Mutagenesis Kit (New England BioLabs; E0554S). For *E. coli* expression the NS5A D1 coding sequence (amino acids 35-215) was PCR amplified and cloned into pET-28a-Sumo using BamHI/XhoI restriction sites. Lentivirus constructs for silencing of CypA and CypB were kindly provided by Prof. Greg Towers (UCL) [19]. CypA wildtype and H126Q catalytically inactive mutant cloned in pGEX-6p-2 were described previously [7].

### Antibodies

The following primary antibodies were used: rabbit anti-CypA (Enzo; BML-SA296-0100), rabbit anti-CypB (Abcam;ab16045), mouse-anti β-Actin (Sigma Aldrich; A1978), sheep anti-GST (in-house) and sheep-anti NS5A (in house) [24]. Secondary IRDye 680 and 800 labelled antibodies were obtained from Li-COR.

### Electroporation and luciferase assay

Huh7 and Huh7.5 cells were washed in ice-cold PBS. Cells (5×10^6^) were resuspended in ice-cold PBS and electroporated with 2 µg of RNA at 950 µF, 270 V. Cells were resuspended in complete media and then seeded separately into 96-well plates at 3×10^4^ cells/well. At the indicated times post-electroporation (hpe), cells were harvested into 30 µl passive lysis buffer (PLB; Promega), incubated for 15 min at room temperature and stored at -80°C until assayed. Luciferase activity was assessed (Promega) on a FluoStar Optima luminometer. Data were recorded as relative light units (RLU).

### CsA assay (EC_50_)

Huh7.5 cells were electroporated and seeded as described above in 96-well plates. CsA (Sigma-Aldrich) was resuspended in dimethyl sulfoxide (DMSO) as 20 mM stock, diluted as required and added to cells at 4 hpe. Cells were harvested and luciferase activity determined as above. Data were modelled using the model of log (agonist) vs. response and EC_50_ values were calculated with GraphPad Prism.

### Lentivirus production and construction of stable knockout cell lines

HEK293T cells in 10 cm dishes were transfected with 1 μg packaging plasmid p8.91, 1 μg envelope plasmid pMDG encoding VSV-G protein and 1.5 μg transfer plasmid pHIV-SIREN encoding shRNA to CypA or CypB as described [19]. Lentivirus supernatants were collected at 48 h and filtered through a 0.45 μm syringe. Huh7 or Huh7.5 cells were seeded into 6 well plates at density of 2.5 × 10^5^ cells/well and transduced with 1 ml/well lentivirus and 8 μg/ml polybrene for 24 h. Transduced cells were selected using 2.5 μg/ml puromycin at 72 h post transduction.

### Western blotting

Cells were washed twice in ice-cold PBS and lysed in GLB (1 % Triton X-100, 120 mM KCl, 30 mM NaCl, 5 mM MgCl_2_, 10 % glycerol (v/v), and 10 mM piperazine-N,N’-bis (2-ethanesulfonic acid) (PIPES)-NaOH, pH 7.2) with protease and phosphatase inhibitors (Roche; 5892791001). 20 µg of each sample were denatured at 95°C for 5 min and separated by SDS-PAGE. Proteins were transferred to polyvinylidene fluoride (PVDF) membrane and blocked with 50 % (v/v) Odyssey blocking buffer (Li-COR) diluted in 1x Tris-buffered saline containing Tween-20 (TBS-T) (50 mM Tris-HCl pH 7.4, 150 mM NaCl, 0.1% Tween-20). Membranes were probed with primary antibodies (1:1000 dilution) at 4°C overnight, and stained with IRDye labelled anti-mouse (700 nm) and anti-rabbit (800 nm) secondary antibodies for 1 h at room temperature (RT). Membranes were imaged on a LI-COR Odyssey Sa Imager.

### His-SUMO-NS5A D1 and GST-CypA expression, purification and pulldown assay

Plasmids were freshly transformed into *Escherichia coli* BL21 (DE3) pLysS and grown at 37°C until OD_600_ values reached 0.6-0.8. Protein expression was induced by 100 μM isopropyl β-D-1-thiogalactopyranoside (IPTG) at 18°C for 6 h. Cells were recovered by centrifugation at 8000 × g for 15 min and resuspended in 50 ml binding buffer (100 mM Tris pH 8.2, 200 mM NaCl, 20 mM imidazole), supplemented with 40 μl DNase, 40 μl RNaseA, 2 mg/ml lysozyme and protease inhibitors (Roche) per 1 L of culture. After incubation on ice for 30 min, samples were sonicated: 10 microns for 12 pulses of 20 sec separated by 20 sec at 4°C. After centrifugation at 4000 × g for 1 h at 4°C twice, supernatants were filtered through a 0.45 μm syringe filter. Samples were transferred to 1 ml HisTrap FF or GST-agarose columns (Cytiva) equilibrated with binding buffer and the columns were washed 3 times using 5 column volumes of binding buffer. The samples were eluted with binding buffer containing 250 mM imidazole or GST elution buffer (50 mM Tris-HCl, 10 mM reduced glutathione, pH 8.0) and dialyzed against 20 mM Tris-HCl, pH 8.2, 150 mM NaCl and 10 % (v/v) glycerol for His-SUMO-NS5A DI or 50 mM Tris-HCl, pH 7.5, 100 mM NaCl and 10 % (v/v) glycerol for GST-CypA.

20 µL GST beads was transferred into 1.5 mL Eppendorf tubes. After washing the beads twice with 400 µL binding buffer (100 mM Tris, 0.5 M NaCl, 1 % Triton X100), 5 µg GST (diluted with washing buffer) protein was added to beads and incubated at 4°C for 2 h. Beads were washed 5 times with binding buffer for 5 min and blocked overnight in 400 µL washing buffer with 3 % BSA. 1.5 µg His-Sumo NS5A D1 protein diluted with washing buffer was added and incubated at 4°C for 2 h. After 5 washes using binding buffer, the bound material was eluted with 20 µL SDS sample buffer, heated for 10 min at 95°C and analyzed by western blotting.

### Statistical analysis

Statistical analysis was performed using an unpaired two-tailed Student’s t tests on GraphPad Prism version 9.30. **** (P<0.0001) and *** (P<0.001) indicate significant difference from wild type (n>=3). Data in histograms are displayed as the means ± S.E.

## References

1. Younossi ZM, Wong G, Anstee QM, Henry L. The Global Burden of Liver Disease. Clin Gastroenterol Hepatol 2023;21(8):1978–1991.

2. Chan A, Patel K, Naggie S. Genotype 3 Infection: The Last Stand of Hepatitis C Virus. Drugs 2017;77(2):131–144.

3. Smith D, Magri A, Bonsall D, Ip CLC, Trebes A et al. Resistance analysis of genotype 3 hepatitis C virus indicates subtypes inherently resistant to nonstructural protein 5A inhibitors. Hepatology 2019;69(5):1861–1872.

4. Simmonds P, Becher P, Bukh J, Gould EA, Meyers G et al. ICTV Virus Taxonomy Profile: Flaviviridae. J Gen Virol 2017;98(1):2–3.

5. Yin C, Goonawardane N, Stewart H, Harris M. A role for domain I of the hepatitis C virus NS5A protein in virus assembly. PLoS Pathog 2018;14(1):e1006834.

6. Chen S, Harris M. NS5A domain I antagonises PKR to facilitate the assembly of infectious hepatitis C virus particles. PLoS Pathog 2023;19(2):e1010812.

7. Foster TL, Gallay P, Stonehouse NJ, Harris M. Cyclophilin A interacts with domain II of hepatitis C virus NS5A and stimulates RNA binding in an isomerase-dependent manner. J Virol 2011;85(14):7460–7464.

8. Yang F, Robotham JM, Grise H, Frausto S, Madan V et al. A major determinant of cyclophilin dependence and cyclosporine susceptibility of hepatitis C virus identified by a genetic approach. PLoS Pathog 2010;6(9).

9. Liu Z, Yang F, Robotham JM, Tang H. Critical role of cyclophilin A and its prolyl-peptidyl isomerase activity in the structure and function of the hepatitis C virus replication complex. J Virol 2009;83(13):6554–6565.

10. Verdegem D, Badillo A, Wieruszeski JM, Landrieu I, Leroy A et al. Domain 3 of NS5A protein from the hepatitis C virus has intrinsic alpha-helical propensity and is a substrate of cyclophilin A. J Biol Chem 2011;286(23):20441–20454.

11. Watashi K, Ishii N, Hijikata M, Inoue D, Murata T et al. Cyclophilin B is a functional regulator of hepatitis C virus RNA polymerase. Mol Cell 2005;19(1):111–122.

12. Harding MW, Handschumacher RE. Cyclophilin, a primary molecular target for cyclosporine. Structural and functional implications. Transplantation 1988;46(2 Suppl):29S–35S.

13. Mamatis JE, Pellizzari-Delano IE, Gallardo-Flores CE, Colpitts CC. Emerging Roles of Cyclophilin A in Regulating Viral Cloaking. Front Microbiol 2022;13:828078.

14. Fischer G, Gallay P, Hopkins S. Cyclophilin inhibitors for the treatment of HCV infection. Curr Opin Investig Drugs 2010;11(8):911–918.

15. Targett-Adams P, McLauchlan J. Development and characterization of a transient-replication assay for the genotype 2a hepatitis C virus subgenomic replicon. J Gen Virol 2005;86:3075–3080.

16. Hughes M, Griffin S, Harris M. Domain III of NS5A contributes to both RNA replication and assembly of hepatitis C virus particles. J Gen Virol 2009;90(Pt 6):1329–1334.

17. Ramirez S, Mikkelsen LS, Gottwein JM, Bukh J. Robust HCV Genotype 3a Infectious Cell Culture System Permits Identification of Escape Variants With Resistance to Sofosbuvir. Gastroenterology 2016;151(5):973–985 e972.

18. Ward JC, Bowyer S, Chen S, Fernandes Campos GR, Ramirez S et al. Insights into the unique characteristics of hepatitis C virus genotype 3 revealed by development of a robust sub-genomic DBN3a replicon. J Gen Virol 2020;101(11):1182–1190.

19. Colpitts CC, Ridewood S, Schneiderman B, Warne J, Tabata K et al. Hepatitis C virus exploits cyclophilin A to evade PKR. Elife 2020;9.

20. Hanoulle X, Badillo A, Wieruszeski JM, Verdegem D, Landrieu I et al. Hepatitis C Virus NS5A Protein Is a Substrate for the Peptidyl-prolyl cis/trans Isomerase Activity of Cyclophilins A and B. J Biol Chem 2009;284(20):13589–13601.

21. Sumpter R, Jr., Loo YM, Foy E, Li K, Yoneyama M et al. Regulating intracellular antiviral defense and permissiveness to hepatitis C virus RNA replication through a cellular RNA helicase, RIG-I. J Virol 2005;79(5):2689–2699.

22. Nakabayashi H, Taketa K, Miyano K, Yamane T, Sato J. Growth of human hepatoma cells lines with differentiated functions in chemically defined medium. Cancer Res 1982;42(9):3858–3863.

23. Blight KJ, McKeating JA, Rice CM. Highly permissive cell lines for subgenomic and genomic hepatitis C virus RNA replication. J Virol 2002;76(24):13001–13014.

24. Macdonald A, Crowder K, Street A, McCormick C, Saksela K et al. The Hepatitis C Virus Non-structural NS5A Protein Inhibits Activating Protein–1 Function by Perturbing Ras-ERK Pathway Signaling. Journal of Biological Chemistry 2003;278(20):17775–17784.

