## Supplementary figures S1-S3 for "Mutational analysis reveals a novel role for hepatitis C virus NS5A domain I in cyclophilin-dependent genome replication"

```
       1                                                      50
JFH1   SGSWLRDVWD WVCTILTDFK NWLTSKLFPK LPGLPFISCQ KGYKGVWAGT
DBN3a  SGDWLRDIWD WVCTVLSDFK SWLSAKIMPA LPGLPFISCQ KGYKGVWRGD
Cons   SG.WLRD.WD WVCT.L.DFK .WL..K..P. LPGLPFISCQ KGYKGVW.G.

       51                                                    100
JFH1   GIMTTRCPCG ANISGNVRLG SMRITGPKTC MNTWQGTFPI NCYTEGQCAP
DBN3a  GVMSTRCPCG ATIAGHVKNG SMRLAGPRTC ANMWYGTFPI NEYTTGPSTP
Cons   G.M.TRCPCG A.I.G.V..G SMRL.GPKTC .N.W.GTFPI N.YT.G...P

       101                                                   150
JFH1   KPPTNYKTAI WRVAASEYAE VTQHGSYSYV TGLTTDNLKI PCQLPSPEFF
DBN3a  CPSPNYTRAL WRVAASSYVE VRRVGDFHYI TGATEDELKC PCQVPAAEFF
Cons   .P..NY..A. WRVAAS.Y.E V...G...Y. TG.T.D.LK  PCQ.P..EFF

       151                                                   200
JFH1   SWVDGVQIHR FAPTPKPFFR DEVSFCVGLN SYAVGSQLPC EPEPDADVLR
DBN3a  TEVDGVRLHR YAPPCKPLLR DDITFMVGLN SYAIGSQLPC EPEPDVSVLT
Cons   ..VDGV..HR .AP..KP..R D...F.VGLN SYA.GSQLPC EPEPD..VL.

       201                                                   250
JFH1   SMLTDPPHIT AETAARRLAR GSPPSEASSS VSQLSAPSLR ATCTTHSNTY
DBN3a  SMLRDPSHIT AETAARRLAR GSPPSEASSS ASQLSAPSLK ATCQTHRPHP
Cons   SML.DPSHIT AETAARRLAR GSPPSEASSS .SQLSAPSL. ATC.TH....

       251                                                   300
JFH1   DVDMVDANLL MEGGVAQTEP ESRVPV---- LDFLEPMAEE ESDLEPSIPS
DBN3a  DAELVDANLL WRQEMGSNIT RVESETKVVI LDSFEPLRAE IDDAELSVAA
Cons   D...VDANLL .......... ......---- LD..EP...E ..D.E.S...

       301                                                   350
JFH1   ECMLPRSGFP RALPAWARPD YNPPLVESWR RPDYQPPTVA GCALPPPKKA
DBN3a  ECFKKPPKYP PALPIWARPD YNPPLLDRWK APDYEPPTVH GCALPPRGAP
Cons   EC........ .ALP.WARPD YNPPL...W. .PDY.PPTV. GCALPP....

       351                                                   400
JFH1   PTPPPRRRRT VGLSESTISE ALQQLAIKTF GQPPSSGDAG SSTGAGAAES
DBN3a  PVPPPRRKKT IQLDGSNVSA ALAALAEKSF PSSKPQEENS SSSGVDTQSS
Cons   P.PPPRR..T ..L..S..S. AL..LA.K.F .......... SS.G.....S

       401                                      ▼            450
 JFH1  GGPTSPGEPA PSETGSA.SS MPPLEGEPGD PDLESDQVEL QPPPQGGGVA
DBN3a  ITSKVPPSPG GESDSESCSS MPPLEGEPGD PDLSCD---- ----------
Cons   .....P..P. ........SS MPPLEGEPGD PDL..D---- ----------

       451                  472
JFH1   PGSGSGSWST CS--EE.DDTTV CC
DBN3a  ------SWST VSDSEE--QSVV CC
Cons   ------SWST .S——EE--...V CC
```

Supplementary Figure S1. Alignment of JFH-1 and DBN3a NS5A amino acid sequence. Cons: consensus identity. Dashes indicate where the sequences do not align and represent insertions. Residues bolded were subject to alanine scanning mutagenesis in this study.

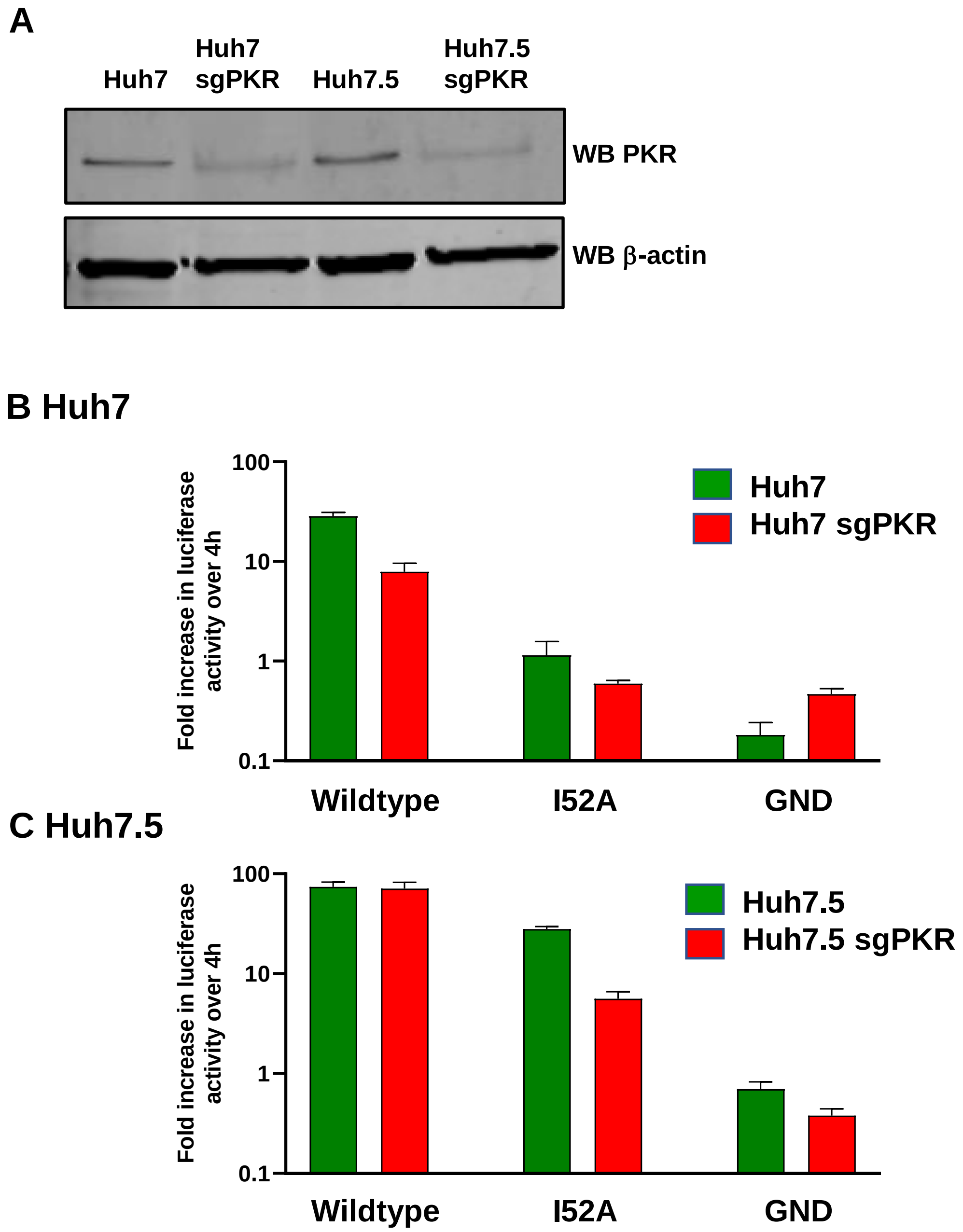


**Supplementary Figure S2. SGR-luc-JFH-1 replication in PKR-silenced cells. A** WB of control or PKR silenced cells. **B, C** The indicated cells were electroporated with SGR-luc-JFH-1 wildtype or I52A mutant RNA and harvested at 48 hpe. Firefly luciferase activity was normalized with respect to 4 hpe.

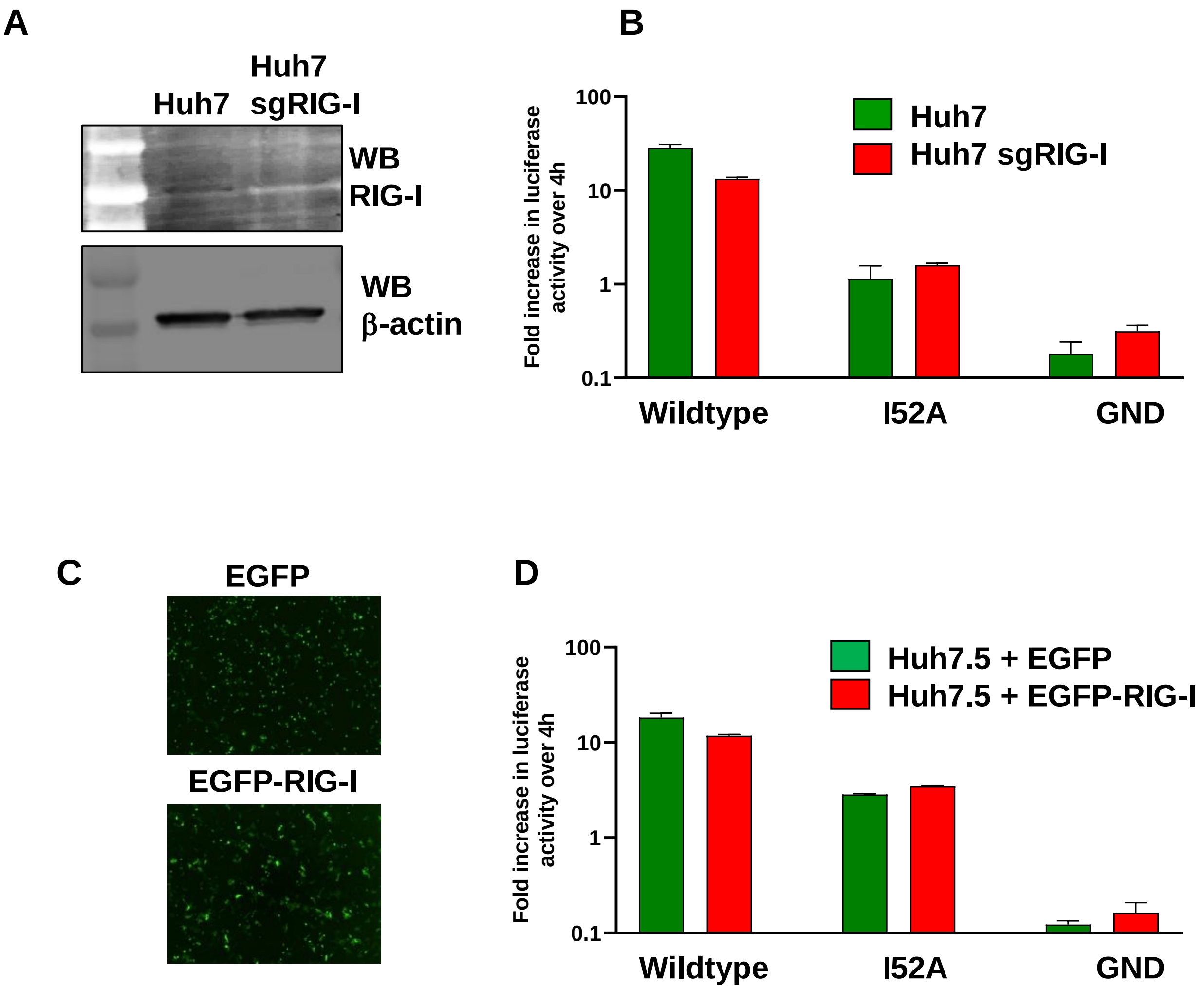


**Supplementary Figure S3. SGR-luc-JFH-1 replication is not affected by RIG-I expression.** **A** WB of control or RIG-I silenced Huh7 cells. **B** The indicated cells were electroporated with SGR-luc-JFH-1 wildtype or I52A mutant RNA and harvested at 48 hpe. Firefly luciferase activity was normalized with respect to 4 hpe. **C** Fluorescence microscopy of Huh7.5 cells transfected with pEGFP or pEGFP-RIG-I. **D** Huh7.5 cells transfected with pEGFP or pEGFP-RIG-I were electroporated with SGR-luc-JFH-1 wildtype or I52A mutant RNA and harvested at 48 hpe. Firefly luciferase activity was normalized with respect to 4 hpe.
